# DISCOVERY AND BIOCHEMICAL CHARACTERIZATION OF A FUNGAL ICE NUCLEATION PROTEIN FROM *PODILA CLONOCYSTIS*

**DOI:** 10.64898/2025.12.15.694473

**Authors:** Brenna J. C. Walsh, Sarah L. Brewer, Andrea J. Shepard, Noor Mahmoud, Elizabeth A. Robinson, Brooke M. Luisi, Joel M. Sarapas, William D. Stone

**Affiliations:** Research and Exploratory Development Department, The Johns Hopkins University Applied Physics Laboratory, 11100 Johns Hopkins Road, Laurel, Maryland 20723, USA

## Abstract

Biological ice nucleation plays a pivotal role in atmospheric processes, yet the molecular basis of fungal ice nucleation remains poorly understood compared to bacterial systems. Here, we report the biochemical characterization of an ice nucleation protein (*Pc*INP) from a soil-dwelling fungus *Podila clonocystis,* not previously reported to produce ice nuclei. Using sequence similarity network analysis, we identified *Pc*INP as a putative fungal homolog of bacterial ice nucleation proteins and confirmed its function through recombinant expression in *Escherichia coli.* We probe the function of *Pc*INP structure through domain truncations and demonstrate that a poorly structured N-terminal region is not necessary for ice nucleation activity and can be functionally replaced with an expression enhancing SUMO fusion tag. Finally, we observe both monomeric and aggregated *Pc*INP in *E. coli* lysates using SEC-MALS but are unable to distinguish their ice nucleation activity pointing to an unknown *in vitro* aggregation mechanism. Our findings establish *Pc*INP within the emerging class fungal ice nucleation protein with distinct structural features and high stability, expanding the known diversity of biological ice nucleators and highlighting their potential for environmental and biotechnological applications.

## INTRODUCTION

Supercooling is a thermodynamic phenomenon in which a liquid remains unfrozen at or below its theoretical freezing point. In the case of pure water ice formation is thermodynamically favored below 0 °C, yet is often delayed, with supercooling persisting down to −42 °C due to the high activation barrier to homogenous seed crystal (nuclei) formation.^1^ In contrast, impure water freezes more readily due to heterogeneous nucleation such as the presence of suspended particles and/or macromolecules that serve as ice nuclei to catalyze crystal formation. While many types of nuclei exist, several biological organisms including bacteria, fungi, plants, and lichen that are abundant in the Earth’s atmosphere have been shown to produce them (termed biological ice nuclei). The abundance of these biological ice nuclei in the Earth’s atmosphere are thought to be a major driver of freezing processes above −15 °C that play a role in climatic processes.^2–6^

The best characterized biological ice nuclei are produced by pathogenic, plant-associated gammaproteobacteria such as *Pseudomonas*, *Pantoea*, and *Xanthomonas.* These nuclei are the result of ice nucleation proteins (INPs) that are anchored to the bacterial outer membrane to promote frost damage by nucleating ice crystals, thereby increasing plant susceptibility to infection.^4^ Owing to their activity at relatively high temperatures (up to −2 °C), INPs have also been studied as replacements for mineral ice nuclei like silver iodide (*T_n_* = −6 °C) and Fieldspar (*T_n_ =* −15 °C) investigated for cloud seeding applications.^7,8^ The archetypal, bacterial INP is InaZ from *Pseudomonas syringae*,^9^ which is comprised of three distinct domains: an N-terminal membrane-integrated region tethers the protein to outer membrane,^10^ a central domain of up to 100 tandem 16-residue repeats forming a β-barrel fold with conserved threonine-x-threonine (TxT) and serine-leucine-threonine (SLT) motifs that organize water molecules into clathrate structures that resembles ice-crystal lattices, providing a template for heterogenous nucleation.^11^ Finally, a C-terminal β-barrel extension (∼ 10 repeats) with a short unstructured region is thought to promote the aggregation of multiple INPs into functional units.^11^ Importantly, classical nucleation theory demonstrates that the activity level (*i.e.,* nucleation temperature) of ice nuclei is dependent on the size of their water structuring features and the surface area of their ice-like bound clatharate. INP activity therefore correlates with the total surface area and orientation of the water structuring domains as well as their ability to aggregate into higher-order structures.^12–14^

Bacterial INPs have found niche commercial applications in snowmaking and ice skating rinks, but their widespread use as frost manipulating coatings, biomedical cryopreservatives, and bulk ice additives has been hampered by their poor stability and difficulty of production. A key contributing factr to both of these challenges is bacterial INPs’ membrane mediated aggregation mechanism. The large protein–membrane aggregates that demonstrate high nucleation temperatures (−2 °C) are highly dependent on membrane fluidity and are sensitive to temperatures above 4 °C, pH changes, and ionic strength. These aggregates require cold storage after processing to avoid irreversible denatureation as they can currently only form *in vivo.* Efforts to enhance INP yields through recombinant expression in microbial systems and to purify INPs, for increased potency and reduced non-specific molecule contamination, have failed to maintain the structure of these high activity aggregates.^14^ To mitigate the limitations of bacterial INPs for biotechnology applications, recent efforts has focused on identifying novel biological ice nuclei in the hopes of identifying a more stable aggregation mechanism that can be recapitulated in recombinant or *in vitro* systems.

By far the strongest evidence outside of gram-negative bacteria comes from fungi, with diverse species including *Fusarium, Mortierella, Isaria, Puccinia*, and *Sarocladium* shown to produce proteinaceous ice nuclei.^15–18^ Given the widespread distribution of soil dwelling fungi and the dispersal of their spores as aerosols, fungal INPs are hypothesized to be major contributors to biogenic atmospheric ice nuclei.^19^ Efforts to pinpoint the molecular basis for their ice nucleation activity have focused on *Fusarium*^15^ and *Mortierella.*^18^ Studies in these genera suggest the existence of two distinct classes fungal ice nucleation proteins that differ in size and temperature sensitivity. Both are water soluble, can be washed from the mycelial surface, nucleate between −10 °C and −5 °C, and are sensitive to proteolytic digestion. Filtration studies indicate that *Mortierella* INPs are between 100 and 300 kDa in size, whereas some *Fusarium* INPs readily pass through 100 kDa filters.^15,18^ Heat sensitivity further distinguishes them with certain *Mortieriella* isolates producing ice nuclei stable to heating to 60 °C while *Fusarium* nuclei have reduced activity upon heating to temperatures as low as 40 °C.^15,18^ One study isolated a 5.4 kDa peptide from *Fusarium* using ice affinity chromatography that self-assembled into aggregates of more than 100 monomers, though its exact sequence remains unknown.^20^ More recently, a class of INPs in the *Mortierellaceae* family was identified that closely resembles bacterial INPs, with proteins from *Mortierella alpina* and *Entomortierella parvispora* confirmed in a recombinant yeast system.^21^

Here, we characterized the ice nucleation activity of the fungus *Podila clonocystis,* a member of the *Mortierellaceae* family, and identified its INP using a sequence similarity network approach. We recombinantly expressed this INP in *E. coli,* verified its ice nucleation activity and interrogated a structural model of its domain organization through recombinantly produced truncations. We observed the N-terminal region is not necessary for activity and can be replaced by expression enhancing tags that result in a recombinant protein with enhanced ice nucleation activity and increased the number of active nuclei produced in solution. Notably, *Pc*INP exhibited remarkable thermal stability, making it highly amendable to environmental applications. Finally, we interrogated the aggregation state of this protein using size exclusion chromatography coupled with a multi-angle light scattering detector (SEC-MALS) observing both monomeric and aggregated *Pc*INP in soluble protein lysates. Our findings suggest that *Pc*INP possess a novel mechanism for protein aggregation that may prove durable and more amenable for high yielding expression systems than the membrane mediated mechanism. This work positions *Pc*INP within the emerging class of fungal ice nucleation proteins characterized by distinct structural features and remarkable stability, broadening the landscape of known biological ice nucleators and their potential for environmental and biotechnological applications.

## RESULTS

### Identification of fungal ice nucleation protein

To uncover previously unannotated INPs in public sequence databases, we utilized sequence similarity network (SSN) analysis.^22,23^ SSNs are graphical tools used to visualize relationships between protein sequences based on pairwise similarity, enabling functional and evolutionary analysis across protein families. Using InaZ from *Pseudomonas syringae* as the seed sequence, we generated a network of putative INPs from the UniProt database.^24^ The resulting network comprised 485 sequences, including several previously characterized bacterial INPs (*e.g., Pseudomonas*, *Xanthomonas*, and *Panteoa* species).^9,25,26^ At low alignment thresholds this network formed a single cluster indicative of the highly conserved nature of these proteins. As the alignment score threshold was increased, the majority of sequences remained in a primary cluster containing the InaZ input sequence, but several nodes began to separate (Fig. 1a). These outlying nodes consisted largely of incomplete sequence fragments or sequences from eukaryotic organisms such as fungi, plants, and fish. One node of particular interest corresponded to a putative INP from *Mortierella alpina*, a fungus that was previously shown to have ice nucleation activity.^18^ This sequence is annotated as a fragment on UniProt (A0A9P6LVP5) and is found at the end of a genome scaffold.^27^ To obtain a complete sequence, we performed a basic local alignment search, which identified a homologous protein in the closely related fungus *Podila clonocystis* (Supplementary Fig. 1).^27^ This putative INP was absent from the SSN because protein annotations for *P. clonocystis* are yet included in the UniProt database. Eufemio et al. recently identified and expressed *Mortierella* and *Entomoreriella* INPs in a recombinant yeast system and while they identified a similar *Pc*INP sequence, this sequence has not been experimentally investigated.^25^

**Fig 1.**
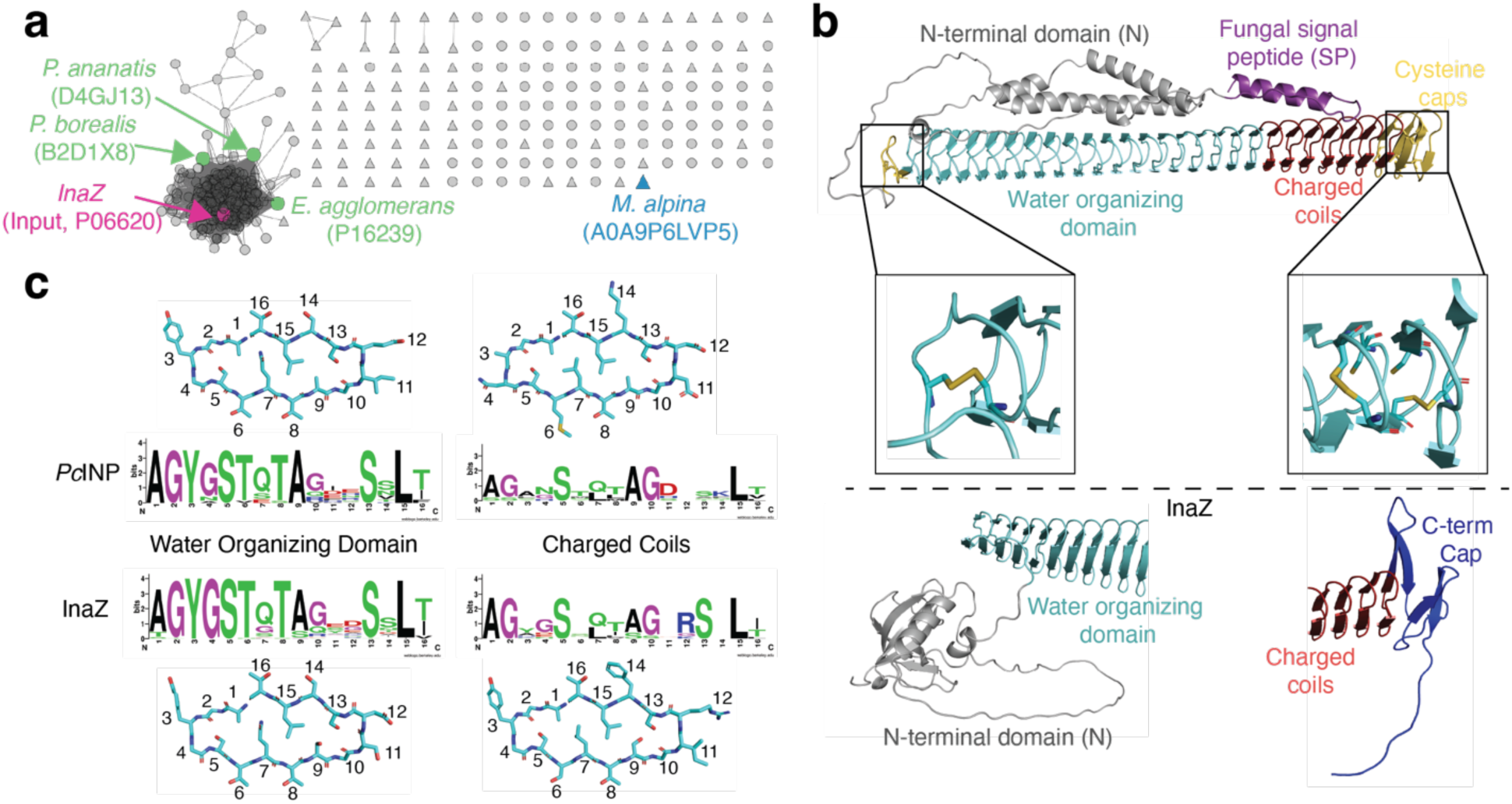
Identification and structural analysis of putative *P. clonocystis* INP. **a**, INP SSN generated using *P. syringae* InaZ as the seed sequence shown in pink. Green nodes indicate additional bacterial INPs that have been characterized. Triangle nodes represent eukaryotic proteins, and putative *M. alpina* INP is shown in blue. **b**, Structural model of *Pc*INP colored according to predicted functional domains. Comparison of *Pc*INP domains to N-terminal and C-terminal domains of InaZ below the dashed line. **c**, Repeat peptide comparison between *Pc*INP and bacterial InaZ for water organizing domain and charged coil repeats.

### Structural prediction of fungal ice nucleation protein

To investigate the domain structure of this putative fungal INP (termed *Pc*INP hereafter), we generated a predicted structural model using Chai Discovery^28,29^ and interrogated the sequence with functional domain prediction tools including SignalP and Phobius (Fig. 1b and Supplementary Fig. 1). The predicted protein architecture included a signal peptide, a N-terminal domain, a central β-barrel repeating domain, and cysteine-rich caps (Fig. 1b). Similar to bacterial INPs, *Pc*INP contains a central β-barrel domain composed of tandem highly conserved 16-amino acid repeating units. These repeats are divided into two regions: a water-structurng domain and charged coils. *Pc*INP encodes 18 repeats of the water organizing domain containing TxT and SLT motifs that form hydrogen-bonded water clathrates. Immediately downstream are 7 charged coil repeats which do not strictly maintain water organizing motifs and contain more charged residues. It should be noted that *Pc*INP contains significantly fewer repeating regions than those of bacterial INPs, which contain over 50 repeats of the water organizing domain and typically 10-12 charged coil repeats. Nevertheless, the water-structuring motifs are highly conserved between *Pc*INP and InaZ (Fig. 1c). The charged coils in *Pc*INP are less conserved and feature a negatively charged aspartic acid in position 11 and a positively charged lysine in position 14 whereas InaZ shows conservations of arginine and serine in positions 12 and 13 (Fig. 1c). Amongst bacterial INPs there is remarkable conservation in the number of these charged coil repeats (10-12) and they been shown to be critical in the multimerization of INP monomers into megadalton sized functional units and ice nucleation activity (> –20 °C).^30^ A lack of conservation in these charged coiled domains suggests *Pc*INP may possess a different mechanism for aggregation.

Another notable difference in *Pc*INP is the presence of cysteine-rich caps flanking the β-barrel.^21^ The N-terminal end of *Pc*INP’s β-barrel domain contains two cysteines, while the C-terminus contains six. Although Chai Discovery does not explicitly predict disulfide bonds, its training on observed folding patterns can generate structural models consistent with disulfide formation. Within multiple independent structure predictions four C-terminal cysteines and the two N-terminal cysteines form intramolecular disulfide bonds within two modified loops of the β-barrel structure (Fig. 1b and Supplemental Fig. 2). The remaining two cysteines at the C-terminus are 4 Å apart but in orientations not conducive to intramolecular disulfide bonding, leaving open the possibility of intermoleculear disulfide linkage. By comparison, InaZ contains four cysteines (three in the N-terminal domain and one in the C-terminal domain) that have not been well studied and are spatially separated from the β-barrel structure. In addition, the InaZ C-terminal domain, shown to be essential for ice nucleation activity, is comparatively longer and more distinct from the β-barrel structure compared to *Pc*INP (Fig. 1b).^31^

A final key difference lies in the N-terminal domains of InaZ and *Pc*INP (Fig. 1b). Both Phobius and SignalP predict the first 20 residues of *Pc*INP encode a secretion signal sequence (Supplementary Fig. 1). The remainder of the N-terminal domain was poorly predicted by our Chai Discovery model but is likely to form a series of ɑ-helices as observed in multiple independent predicted structures (Supplementary Fig. 2). Unlike bacterial INPs, which are membrane anchored, the presence of a signal peptide and absence of transmembrane domains suggest that *Pc*INP is likely secreted from the fungus without membrane association. This is consistent with the observation that fungal INPs readily wash away from biomass and pass through 0.22 μm filters.^15,18^

### Confirmation of *P. clonocystis* ice nucleation activity

Since *P. clonocystis* has not previously been reported to produce ice nuclei, we sought to confirm the ice nucleation activity of this fungus and validate its putative INP in a recombinant system. We obtained two strains of *P. clonocystis* (CBS 176.74 and CBS 357.76) to measure the ice nucleation activity of the native fungus and generated plasmids for recombinant expression of the full-length protein in *E. coli*. Using a microwell plate-based ice nucleation assay (Fig. 2), we demonstrated for the first time that that *P. clonocystis* exhibits ice nucleation activity. Both fungal strains showed similar activity, nucleating at approximately –7 to –8 °C when mycelium was washed with buffer and filtered (0.22 μm) indicating that ice nuclei were not cell associated. Suspensions of intact *E. coli* cells expressing recombinant full-length *Pc*INP constructs, which differed by the presence or absence of C-terminal HA epitope and hexahistidine tags, were also tested. These constructs nucleated ice around –9 °C, slightly below the activity of natively produced *P. clonocystis* ice nuclei. The equivalent nucleation activity of both constructs indicates that C-terminal tags do not affect *Pc*INP function; therefore, the tagged construct was used in all subsequent experiments.

**Fig 2.**
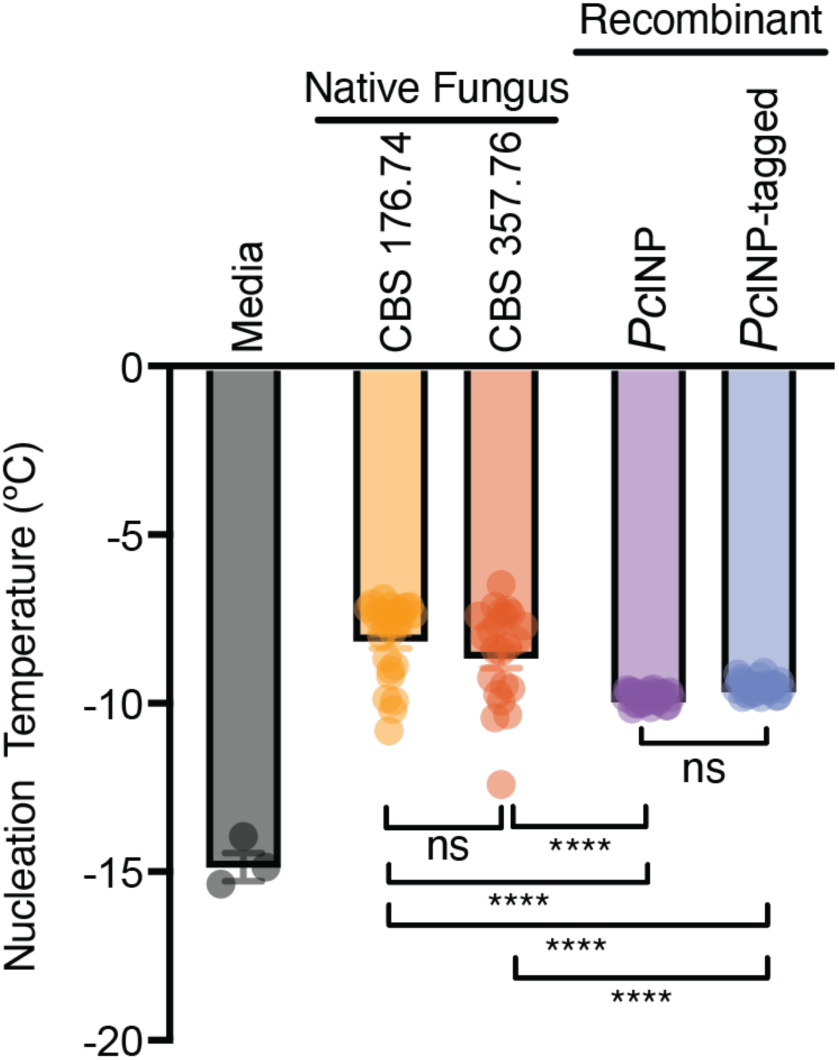
Ice nucleation activity of *P. clonocystis* and recombinant *Pc*INP. Nucleation temperature for two strains of *P. clonocystis* (CBS 176.74 and CBS 357.76) and *E. coli* cells recombinantly expressing *Pc*INP with and without peptide tags for affinity purification. Data are presented as mean ± s.e.m. Statistical analysis was performed using a one-way ANOVA (**** *p ≤* 0.0001, n.s. *p ≥* 0.05). All samples have *p*-value less than or equal to 0.0001 relative to media control.

### Biochemical characterization of INP from *P. clonocystis*

To further probe the roles of different *Pc*INP domains in ice nucleation, we generated a series of truncation and fusion variants in this bacterial recombinant system (Fig. 3). Constructs included: (i) removal of the signal peptide (*Pc*INP-ΔSP), (ii) further deletion of the N-terminal domain (residues 2-166; *Pc*INP-ΔSPΔN), and (iii) addition of an N-terminal small ubiquitin-like modifier (SUMO)^32^ tag to improve expression and solubility levels (SUMO-*Pc*INP-ΔSP and SUMO-*Pc*INP-ΔSPΔN; Supplementary Table 1).

**Fig 3.**
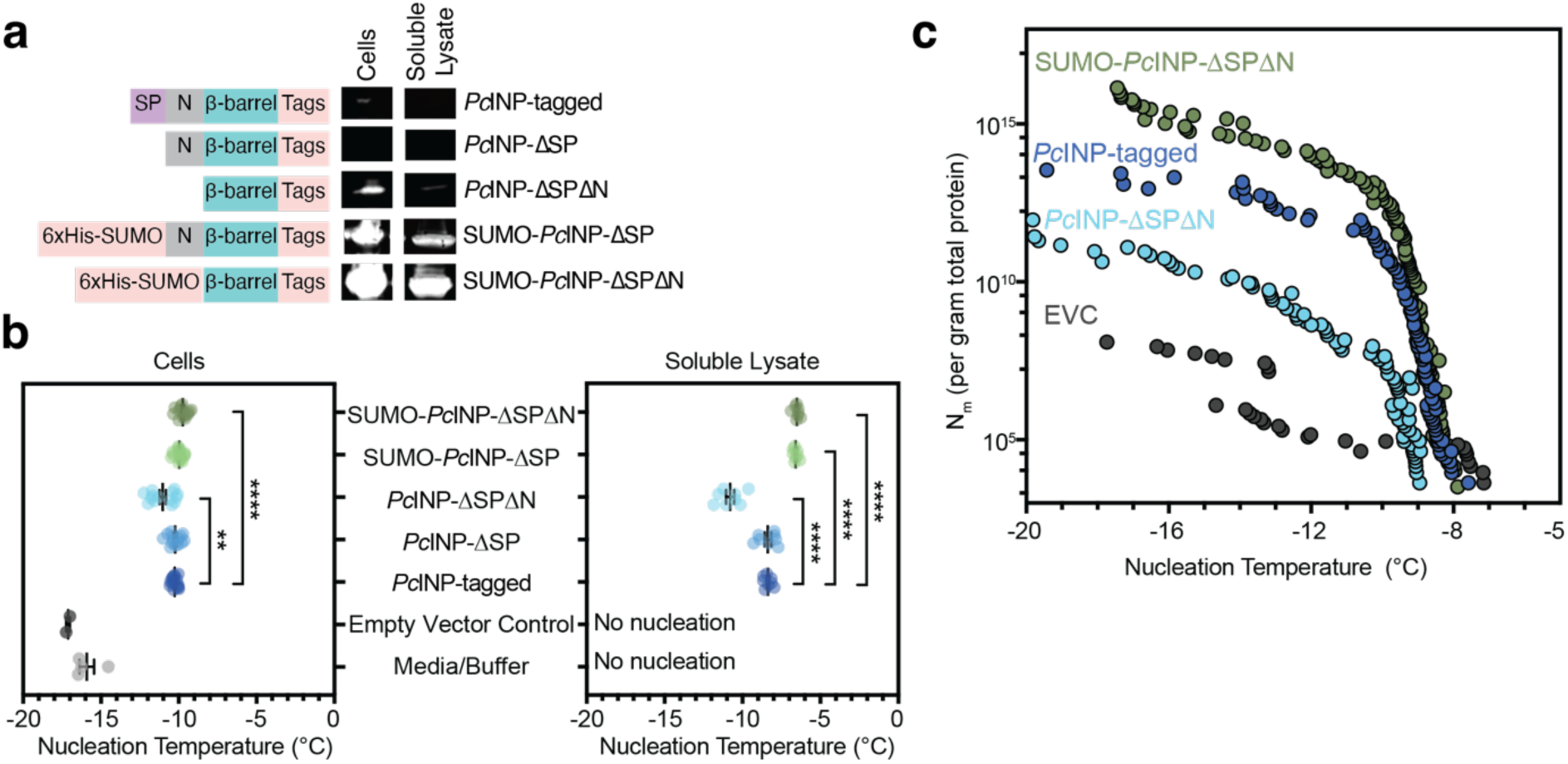
Biochemical characterization of *Pc*INP. **a**, Anti-HA western blot analysis of *E. coli* cells or soluble protein lysates of *Pc*INP variants. Diagrams depicting domains of each variant are shown for each. C-terminal tags include a TEV cleavage site, HA epitope, and hexahistidine tag except for SUMO tagged variants where the hexahistidine tag is N-terminal. **b**, Ice nucleation activity measured on *E. coli* cells (left) or soluble protein lysates (right). Samples were concentration normalized based on OD_600_ for cells and total soluble protein concentration for soluble lysates. Data are presented as mean ± s.e.m. Statistical analysis was performed using a one-way ANOVA (**** *p ≤* 0.0001, ** *p ≤* 0.01). Samples with statistically significant changes relative to *Pc*INP-tagged are included. All samples have *p*-value less than or equal to 0.0001 relative to media, buffer, and empty vector controls. **c**, Number of ice nuclei (N_m_) per gram of total soluble protein measured for soluble protein lysates of select *Pc*INP variants and empty vector control (EVC).

Protein expression and solubility were assessed *via* western blotting of total cellular protein and the soluble protein fraction (Fig. 3a). The full length protein (*Pc*INP-tagged) and the variant without the putative signal peptide (*Pc*INP-ΔSP) displayed poor expression and solubility in this recombinant system. Truncation of the N-terminal domain (*Pc*INP-ΔSPΔN) improved the expression of the protein but it remained poorly soluble. In contrast, addition of a SUMO tag improved the expression and solubility of the recombinant protein both with (SUMO-*Pc*INP-ΔSP) and without the N-terminal domain (SUMO-*Pc*INP-ΔSPΔN), with the later construct exhibiting the highest degree of expression and solubility in a qualitative assessment.

We next investigated how these recombinant INPs performed in an ice nucleation assay by measuring both resuspended whole cells and post-lysis soluble protein fractions (Fig. 3b). We found that all variants displayed ice nucleation activity in whole cells between –9 to –11 °C and were significantly higher that both media and empty vector controls (EVC), indicating despite being near the detection limit by western blot, sufficient protein was present to observe ice nucleation activity. While most variants displayed similar activities, the nucleation temperature of *Pc*INP-ΔSPΔN demonstrated a statistically significant decrease relative to *Pc*INP-tagged that was reversed by the addition of the SUMO tag. Interestingly, more robust trends became apparent when ice nucleation activity was measured on the soluble protein fraction of cellular lysates. While it is impossible to directly compare INP concentrations between whole cells and soluble fractions, all soluble fractions nucleated at higher temperatures (–6 to –11 °C) than in whole cells. This is likely due to a lower solute concentration compared to the cytosol in whole cells leading to a reduction in the colligative depression of freezing temperature. When compared to *Pc*INP-tagged, which nucleated around –8 °C, deletion of the signal peptide alone did not impact activity but further truncation of the N-terminal domain had a statistically significant decrease in nucleation temperature (around –11 °C). The SUMO tagged variants conversely had statistically significant increase in activity and nucleated around –6.5 °C regardless of whether or not the N-terminal domain was present.

We hypothesized the difference in activity of between the full length (*Pc*INP-tagged) and N-terminal trucation *Pc*INP (*Pc*INP-ΔSPΔN) was due to changes in the concentrations of functional INPs present in these recombinant systems. To test this, we performed a dilution series ice nucleation assay on the soluble protein fraction of three variants (*Pc*INP-tagged, *Pc*INP-ΔSPΔN, and SUMO-*Pc*INP-ΔSPΔN) and the EVC starting at 1 mg/mL total protein and diluting until ice nuclei were exhausted (Fig. 3c). In doing so we found that all three variants qualitatively exhibit a monotonic increase to a single plateu in contrast to a multiphasic curve suggesting recombinantly produced *Pc*INP contains a single class of ice nuclei. In addition, the trends in the maximum nucleation temperature were consistent with the number of nuclei present and followed the trend SUMO-*Pc*INP-ΔSPΔN > *Pc*INP-tagged > *Pc*INP-ΔSPΔN (Fig. 3).

### Probing the stability of *Pc*INP to redox state and temperature

To determine whether the oxidation state of the cysteine caps impacted INP activity, we pre-treated soluble lysates of *Pc*INP variants with chemical treatments to alter the state of the cysteines including a reducing agent and hydrogen peroxide (Supplementary Fig. 3). Importantly, control samples of buffer, EVC lysates, and Snomax^®^ were tested alongside the variants, all of which received the same pre-treatments. We found that all treatments decreased the nucleation temperature of Snomax^®^ whereas only the reducing agent alone impacted the *Pc*INP variants. Interestingly, variants of *Pc*INP lacking the N-terminal domain exhibited a larger TCEP dependent decrease in nucleation temperature than the full length variant of *Pc*INP, suggesting the N-terminal domain plays a rule in stabilizing the water organizing domain. To further assess stability, we next investigated the impact of heat on *Pc*INP activity. To do this, soluble protein lysates of *Pc*INP variants and Snomax^®^ suspensions were heated to 95 °C and then tested in a dilution series to measure the number of active nuclei (Supplementary Fig. 4). At high concentrations of heat-treated *Pc*INP and SUMO-*Pc*INP-ΔSPΔN retained maximal nucleation temperatures (–9°C) comparable to the untreated samples. However, heat treatment reduced the total number of active nuclei by approximately 10-fold for *Pc*INP and by approximately 10^3-^fold for SUMO-*Pc*INP-ΔSPΔN. In contrast, when Snomax^®^ was heat treated the maximal ice nucleation temperature was decreased by 6 °C (to –9 °C) accompanied by an approximately 10^5^-fold decrease in the number of nuclei. This suggests that *Pc*INP is more heatstable than bacterial INPs and better able to retain ice nucleation activity even when not highly aggregated.

### Measurement of recombinant ice nucleation protein size

Given that potent ice nucleation activity is linked to protein oligomerization in bacterial INPs, we hypothesized that *Pc*INP activity may also depend on its size and aggregation state. To test this, we investigated the size of *Pc*INP using size exclusion chromatography (SEC) coupled to multi-angle light scattering detector (MALS). This approach separates proteins by their hydrodynamic volume and measures their mass and radius of gyration by MALS to correlate protein activity to functional size. Using a series of protein standards we expect high molecular weight species, including aggregated protein, to elute before 30 min and proteins around 40-80 kDa should elute within 40-50 min (Supplementary Fig. 5). The anticipated monomer, dimer, and trimer sizes of SUMO-*Pc*INP-ΔSPΔN are 60.7, 121.5, and 182.2 kDa respectively indicating monomeric species should elute within 40-50 min while INP multimers will elute earlier. Soluble protein lysates of EVC and SUMO-*Pc*INP-ΔSPΔN were separated by SEC-MALS, yielding chromatograms with similar peaks (Fig. 4a). Four UV absorbance peaks monitoring protein content at 280 nm were observed at approximately 26, 46, 66, and 85 min and aligned well between the samples. Based on the protein standards the peak at 26 min likely contains highly aggregated species or protein multimers, 46 min contains proteins in the 40-80 kDa range, and peaks after 55 minutes are likely peptides and other very small species (Supplementary Fig. 5). In addition to absorbance, two light scattering peaks were detected at 25 and 27 min and the intensity of the peaks was different between EVC and INP samples. The calculated masses of these peaks also indicated a mixture of higher molecular weight or aggregated species that cannot be resolved under these conditions.

**Fig 4.**
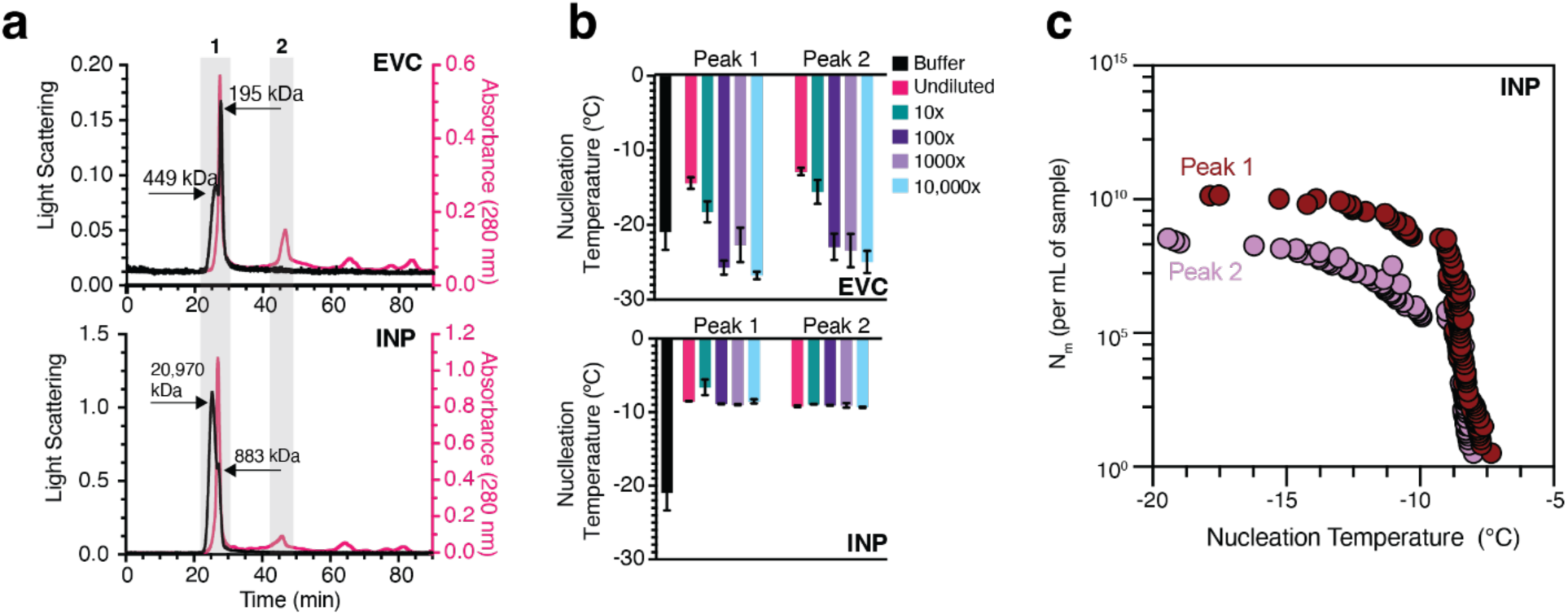
SEC-MALS analysis of *Pc*INP. **a**, Absorbance (280 nm) and light scattering signals for EVC and SUMO-*Pc*INP-ΔSPΔN (INP). **b**, Ice nucleation assay on fractions collected from SEC-MALS and analyzed undiluted and after a series of dilutions. Data are presented as mean ± s.e.m. **c**, Number of ice nuclei (N_m_) per volume of sample measured for SEC-MALS peaks 1 and 2 of the soluble INP lysate.

Based on the retention time of these two peaks, we hypothesized that peak 1 would contain *Pc*INP aggregated as multimers and peak 2 would contain monomeric *Pc*INP. To determine if these peaks contained ice nuclei and to correlate nucleation activity to functional protein size, we collected fractions for these two peaks for both samples and tested them for ice nucleation activity using the plate-based assay at various dilutions (Fig. 4b). Undiluted fractions from the EVC nucleated around –15 °C but decreased to below the buffer control when diluted 100-fold. This INP independent activity is likely due to non-specific nucleation due to high concentrations of protein used to obtain measurable UV absorbance and MALS signals and is expected. In contrast, both peaks of the INP sample demonstrated ice nucleation activity at higher temperatures (around –8 °C) that persisted even with a 10^-4^ dilution indicating that both peaks contained *Pc*INP associated ice nuclei. Importanly, we observed no difference in the maximum nucleation temperature between the monomer and multimer fractions of *Pc*INP. We next sought to quantify the number of nuclei in each peak by performing a full dilution series starting undiluted and diluting until exhaustion (Fig. 4c). These experiments reveal that peak 1 contained an order of magnitude higher concentration of ice nuclei than peak 2 and both peaks exhibit a similar monotonic increase to a single plateau suggesting they contain the same, singular class of ice nuclei.

## DISCUSSION

The identification and characterization of new ice nucleation proteins is key to unraveling biological ice nucleation mechanisms, assessing their influence on biogeochemical cycles, and harnessing them for innovative biotechnologies. Toward the goal of identifying INPs with previously uncharacterized mechanisms of assembly, we used sequence similarity networks to identify putative INPs containing conserved water organizing domains but falling outside of the bacterial INP cluster. Using this approach we identified an uncharacterized INP from *Podila clonocystis*, an organism not previously known to produce ice nuclei. We experimentally confirmed that *P. clonocystis* demonstrates ice nucleation activity and performed a biochemical characterization of its ice nucleation protein, *Pc*INP, in a recombinant system. One unexplained result is the observation that recombinantly produced *Pc*INP has a lower maximum nucleation temperature than extracts prepared from the native fungi (–9 °C vs –7 °C). This reduction in nucleation temperature is similar to what is observed in recombinant bacterial INP expression, where membrane fluidity and an inability to form high activity aggregates is suspected. Further studies may elucidate whether this difference is due to levels of protein expression, interaction with non-peptide fungal components, or host dependent differences in aggregation state.

Utilizing our recombinant production system enabled us to probe the function of domain features unique to fungal INPs. Compared to bacterial INPs, *Pc*INP has fewer repeat regions of the water organizing domain, contains a structurally distinct non-transmembrane N-terminal domain, and cysteine rich caps.^21^ Despite the reduced number of water organizing domains, *Pc*INP retains ice nucleation activity although generally at slightly lower temperatures than longer INPs. One advantage of this smaller size is an increased yield and concentration of nuclei possible due to decreased protein mass. We attempted to further increase yield of *Pc*INP by eliminating extraneous domains by characterizing truncated versions of *Pc*INP. Our results indicate that the N-terminal region is not strictly necessary for *Pc*INP’s ice nucleation activity although deletion of this domain reduced maximum nucleation temperatures indicating it plays some function in modulating activity. The reversal of this decrease in nucleation temperature by the addition of the SUMO solubilizing tag suggests that this maybe due to stabilizing the ice binding domain or modulating overall expression levels rather than modulating aggregation. Another uncharacterized feature of the *Pc*INP structure is the role of two cysteine residues on N-terminal side of the ice binding domain. Our structural model predicted these would form intramolecular disulfide bonds, although we did not experimentally determine this. One hypothesis, is that the N-terminal truncation may have destabilized these cysteine residues. Supporting evidence can be found in the increased susceptibility of N-terminal truncation’s nucleation temperatures to TCEP treatment. Further studies utilizing analytical chemistry methods to detect disulfide bond formation (*e.g.,* mass spectrometry) or spectroscopic methods to probe protein folding such as circular dichroism may illuminate the differences between these constructs. Additionally, genetic engineering of this N-terminal domain may prove a pathforward to enhancing the thermal stability of INPs.

Another notable difference between *Pc*INP and bacterial INPs is in the final charged coiled domains of the β-barrel and the lack of conservation at the C-terminus. Four cysteine residues cap the *Pc*INP C-terminus but only two are predicted to form intramolecular disulfide bonds. The other appear to be oriented to form intermolecular bonds raising the possibility that disulfide bond formation plays a role in aggregation of monomers into multimers or complexation with other molecules. Indeed, it is likely that *Pc*INP has a fundamentally different mechanism of aggregation as it lacks the beta sheets found in the C-terminal aggregation domain of bacterial INPs. Despite the lack of the domain necessary for bacterial INP aggregtes, we directly observed the presence of large aggregates of *Pc*INP in our recombinant lysates but were not able to resolve aggregation state. The observation that there was no difference in the maximum nucleation temperature between the monomer or multimer fractions of *Pc*INP suggests two possibilities. First, that *Pc*INP activity does not depend on aggregation state which is very unlikely (based on nucleation theory and computational models of INP kinetics),^12^ or second that monomers can aggregate into larger protein assemblies *in vitro* after separation by SEC. If proven, this *in vitro* aggregation mechanism would not require biological membranes typically associated with bacterial INP aggregation and would not require a living host to form functional ice nuclei.

While bacterial INPs are commercialized for niche uses, INPs require significant advancement to find broader usage in environmental, material, and biomedical applications. Key barriers to their use in these fields, namely the stability of functional aggregates, yield, and ability to be purified, could be addressed through the further development of recombinant fungal INPs. The thermal induced irreversible activity loss of currently used INP aggregates could be mitigated through the stabilization of INPs using *Pc*INP inspired cysteine caps coupled with an *in vitro* aggregation mechanism. Advances in yield and scalability have already been realized through the use of a recombinant *E. coli* expression system, the smaller size and lack of the membrane domain of *Pc*INP, and the use of the SUMO tag, but could be further enhanced through structural and genetic optimization. Finally, soluble recombinant production and *in vitro* aggregation could allow INPs to be purified by chromatography, increasing their potency and enabling biomedical applications where contamination by host proteins are unacceptable. To fully realize this potential, further investigations should focus on understanding the natural aggegration mechanisms of fungal INPs and engineering *in vitro* assembly of high activity aggreagates before their formulation and use.

## MATERIALS AND METHODS

### Sequence similarity network analysis

The Enzyme Function Initiative Enzyme Similarity Tool (EFI-EST, http://efi.igb.illinois.edu/efi-est/) web tool was used to generate a sequence similarity network (SSN) for ice nucleation proteins.^22,23^ The SSN was generated using the Sequence BLAST option of EFI-EST and the *P. syringae* protein InaZ (P06620) as the seed sequence and default parameters. An alignment score of 100 was used to generate the SSN with no minimum or maximum number of residues, and the final network generated was 100% representative (repnode 100), collapsing sequences of 100% identity over 90% of the sequence into a representative node and was visualized using Cytoscape (http://www.cytoscape.org/).

### Structural model and analysis

Structural model was generated using the Chai Discovery^28,29^ online web tool (https://www.chaidiscovery.com). The results were downloaded and viewed in Pymol. pLogo plots were generated using Weblogo (https://weblogo.berkeley.edu/logo.cgi) and inputs of 16-residue repeats for the β-barrel or charged coil domains of each protein. Plots were generated using the following color scheme green (STNQYC), purple (G), blue (KRH), red (DE), black (AVLIMFWP).

### Cloning of fungal INPs

To recombinantly express the ice nucleation protein from *P. clonocystis* in *E. coli*, the gene encoding BGZ82_000752 (Accession KAG0017472) was codon optimized and cloned into the BamHI and XhoI sites of pET21(+) by Twist Bioscience. A ribosome binding site (TTTGTTTAACTTTAAGAAGGAGA) and NheI restriction enzyme site were inserted upstream of the gene and a SpeI site was added just before the stop codon to make pICE_Podila_001. To make pICE_Podila_002, DNA encoding a TEV protease site, HA tag, and 6x Histidine tag were inserted between the SpeI site and stop codon. For pICE_Podila_005 and pICE_Podila_006, the signal sequence corresponding to amino acids 2-18 of BGZ82_000752 were removed from pICE_Podila_001 and pICE_Podila_002, respectively. For pICE_Podila_009, the N-terminal domain corresponding to amino acids 19-166 were removed from pICE_Podila_006. To increase the expression and solubility of the INP, DNA sequence containing the 6xHis tag was removed from the C-terminus and added with a SUMO tag to the N-terminal of the gene in pICE_Podila_006 and pICE_Podila_009 to make pICE_Podila_007 and pICE_Podila_008 respectively. All genetic components were amplified using Platinum SuperFi II polymerase and assembled using NEBuilder® HiFi DNA Assembly Master Mix. All PCR primers were purchased from Integrated DNA Technologies.

### Recombinant expression of *Pc*INP for assays and purification

Plasmids encoding for recombinant expression of *Pc*INP variants were transformed into *E. coli* BL21 (DE3), cultured in Luria-Bertani (LB) medium at 37 °C with shaking at 250 rpm until the OD_600_ reached 0.6–0.8, induced with 0.5 mM IPTG and expressed at 16 °C for 16-20 h. Two milliliters of cells were transferred to a fresh 2 mL tube and 1 mL was used to measure optical density at OD_600_ using a spectrophotometer. The remaining 1 mL of cells was used in plate-based ice nucleation assays as described below. The remaining cells were harvested by centrifugation (4000 × *g*, 10 min, 25 °C) and either lysed immediately for assays or stored at −80 °C. For ice nucleation assays, the cell pellet was resuspended in a buffer containing 20 mM Tris-HCl pH 7.5, lysed by sonication, then clarified by ultra-centrifugation (119,000 × *g*, 30 min, 4 °C). The resulting soluble protein lysate was quantitated by Bradford assay and used in plate-based ice nucleation assays as described below.

For the purification of SUMO-*Pc*INP-ΔSPΔN, cell pellets were resuspended in lysis buffer containing 25 mM Tris-HCl, 500 mM NaCl, pH 7.5, lysed by sonication and clarified by ultra-centrifugation (119,000 × *g*, 30 min, 4 °C). The soluble protein lysate was filtered through a 0.45 μm syringe filter and loaded onto a 1 mL HisTrap FF Ni-NTA column (Cytiva), which was pre-equilibrated with the same lysis buffer. The column was washed with lysis buffer followed by elution with an imidazole gradient from 0 to 500 mM in the same lysis buffer.

### Westerns

*E. coli* cells grown as specified above were harvested by centrifugation (4000 × *g*, 10 min, 25 °C) and resuspended in 10 mM Tris-HCl pH 7.0 with protease inhibitors (Gold Bio). Samples were sonicated and then clarified in a Sorvall MTX150 micro-ultracentrifuge (ThermoFisher, 117,200 × *g*, 30 min, 4 °C). The supernatant was kept as the soluble protein fraction. Total protein concentration was determined by Bradford assay according to manufacturer’s instructions. Ten μg of protein samples were diluted in Lamelli SDS-Sample buffer and separated on a Novex 4-20% Tris-glycine mini gel. Separated proteins were transferred to a 0.2 μm PVDF membrane using an iBlot 2 Transfer device set to 20 V for 12 min. The membrane was blocked with 5% milk in 1X TBST (150 mM NaCl, 50 mM Tris-HCl, 0.05% Tween-20, pH 7.4) for 1 hr. The membrane was incubated with rabbit antibody against HA tag (Millipore Sigma, H6908) diluted 1:2000 in 1X TBST for 2 hr at room temperature. The membrane was rinsed with fresh 1X TBST three times. The membrane was incubated with an AlexaFluor 488 goat anti-rabbit secondary antibody (Abcam, ab150077) diluted 1:2000 in 1X TBST for 1 hr at room temperature. Proteins were visualized using the 488 channel on an Odyssey M Imaging System (Li-Cor).

### Growth of Podila clonocystis

Two strains of *Podila clonocystis* (CBS 176.74 and CBS 357.76) were received from The Westerdijk Fungal Biodiversity Institute as slant cultures. Mass was transferred via an inoculating loop onto a dextrin-peptone-yeast extract (DPY) plate and incubated at room temperature for at least 7 days for sufficient growth.^18^ Plates were stored at 4 °C until they were assayed. Cells were then collected via gentle scraping into a pre-weighed 15 mL conical tube. 10 mM Tris-HCl buffer pH 7.5 was added to resuspend the fungal material via vortexing to 5 mg/mL. Half the sample was subsequently filtered using a 0.22 μm PES filter. Both the filtered and unfiltered samples were measured in the plate-based ice nucleation assay as described below without further dilution.

### Plate-based ice nucleation assay

To determine the ice nucleation activity, a 384-well plate assay was performed. *E. coli* cell samples were diluted in LB media to an OD_600_ of 0.1 and soluble protein lysates were normalized to a protein concentration of 1×10^-4^ mg/mL in 20 mM Tris-HCl pH 7.5 buffer. Each plate contained a negative control of buffer or media and a positive control of 0.1 mg/mL Snomax^®^. Once diluted, 20 μL of sample was transferred to a 384-well plate in sixteen technical replicates and the plate was briefly centrifuged to ensure samples were at the bottom of each well. The plate is placed inside of an aluminum 384-well block which is then partially submerged in a recirculating bath with either 50% ethylene glycol in water or PolyCool (potassium formate solution, PolyScience). The plate was cooled from 1 to –30 °C over an hour at an approximate rate of 0.5 °C min^-1^. A thermal imaging camera monitors the plate to capture the release of latent heat when a nucleation event occurs upon the well freezing. MATLAB was used to analyze the data from the thermal images to computationally determine the nucleation temperature of each well. To calculate the number of nuclei (N_m_) samples begin at a protein concentration of 1 mg/mL and then diluted in a serial manner 1:10 across multiple orders of magnitude. At least 12 technical replicates per sample were aliquoted into the 384-well plate for testing as described above. The well specific temperatures from MATLAB are further analyzed in Python using a published method.^33^

### SEC-MALS

Soluble protein lysates were separated on a Superdex 200 10/300 GL column (GE Healthcare) using an Agilent 1260 Infinity HPLC. The mobile phase consisted of 20 mM phosphate buffer, 150 mM NaCl, pH 7.4, at a flow rate of 0.5 mL/min. UV absorbance was monitored at 280 nm using a diode-array detector. MALS detection was achieved with a DAWN 8 detector (Wyatt Technology), positioned after the UV detector, and data was processed using ASTRA software (Wyatt Technology). The Debye fitting method was used to analyze the angular dependence of the scattered light and obtain the weight-average molecular weight. Molecular weight determinations were validated by comparing the SEC-MALS results with protein standards to calibrate the system and confirm the reliability of the measurements.

### Statistics and reproducibility

Statistical analyses were conducted using GraphPad Prism v 10.6.1 software (GraphPad Software, LLC). Data are reported as mean ± SEM, unless otherwise specified. Details regarding number of biological and technical replicates are found in the methods. Details regarding the statistical tests applied can be found in the corresponding figure legends.

## ONLINE CONTENT

Any methods, additional references, Nature Portfolio reporting summaries, source data, extended data, supplementary information, acknowledgements, peer review information; details of author contributions and competing interests; and statements of data and code availability are available.

## DATA AVAILABILITY

All data supporting the findings of this study are included in this manuscript. Source data are provided with this paper.

## CODE AVAILABILITY

No custom code was created for this paper.

## ACKNOWLEDGEMENTS

This project has been funded with federal funds from the Defense Advanced Research Projects Agency (DARPA) to J.M.S and W.D.S under Contract No. HR001124F0318.

## AUTHOR CONTRIBUTIONS

B.J.C.W., W.D.S., and J.M.S. supervised the research. B.J.C.W. generated and analyzed SSNs and structural model. S.L.B. and B.M.L. generated plasmids for recombinant protein expression. B.M.L. performed western blots. S.L.B. and A.J.S. performed ice nucleation assays and analyzed data. A.J.S. cultured *P. clonocystis*, measured its activity, and analyzed data. S.L.B. cultured *E. coli* for recombinant protein expression and processed samples for subsequent experiments. N.M was supervised by S.L.B. and A.J.S and assisted for all ice nucleation assays. E.A.R performed all SEC-MALS experiments and analyses. B.J.C.W. wrote the first draft of the manuscript with inputs and edits from all authors.

---

**Supplementary Fig 1.**
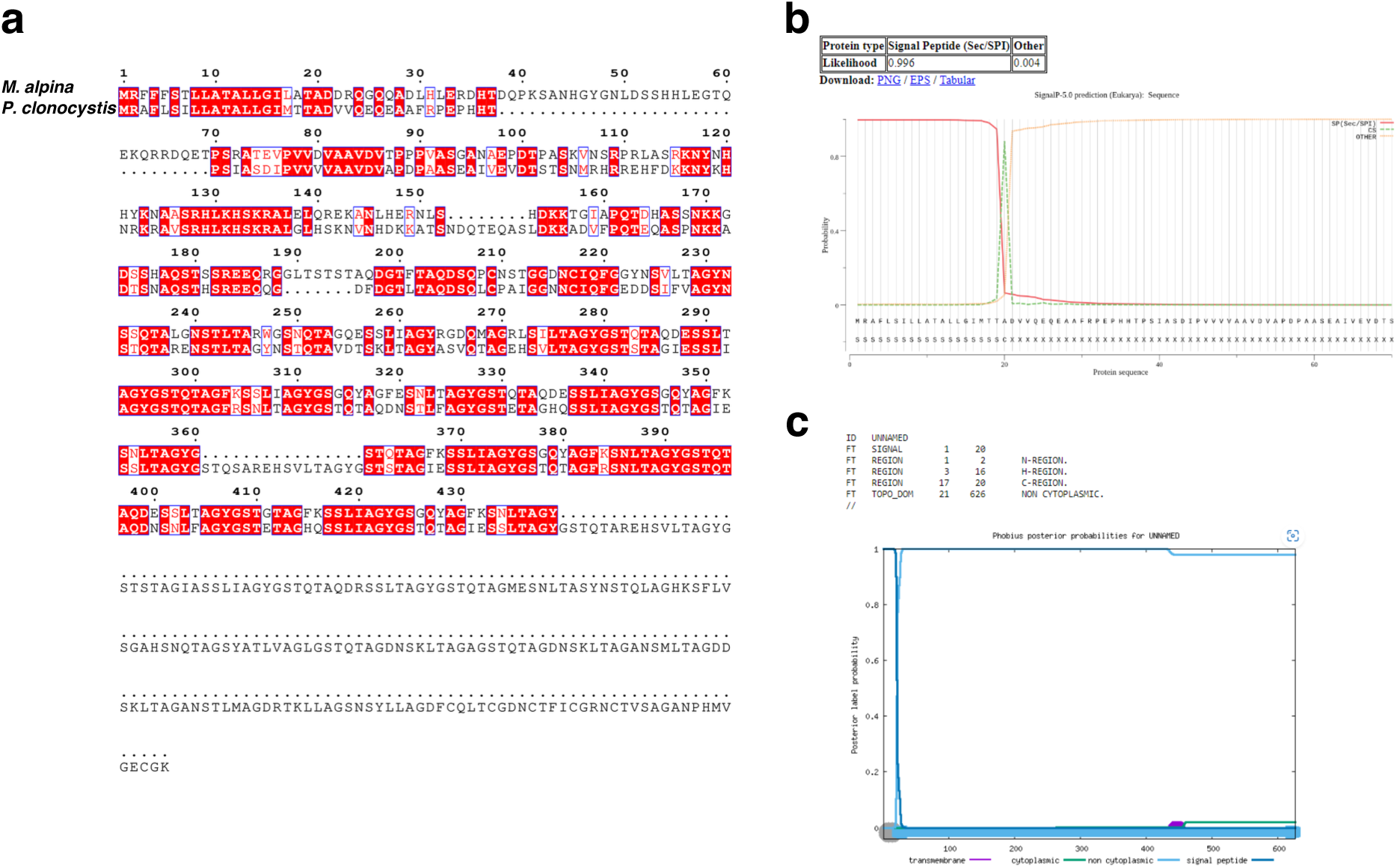
Characterization of putative *P. clonocystis* INP sequence. **a**, Multiple sequence alignment between *M. aplina* (A0A9P6LVP5) and *P. clonocystis* (Accession KAG0017472). **b**, signal peptide prediction from SignalP. **c**, Transmembrane domain prediction from Phobius.

**Supplementary Fig 2.**
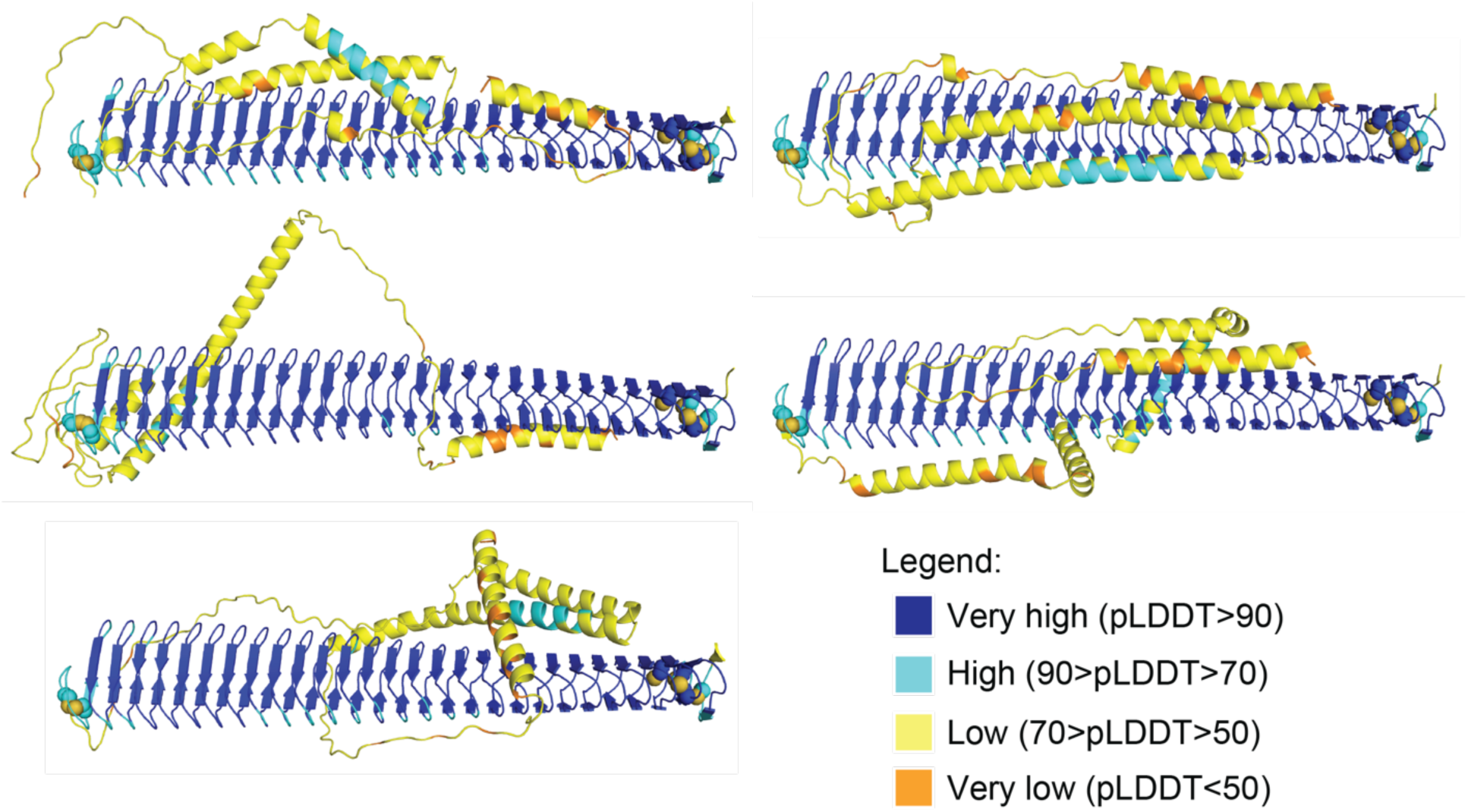
Comparison of predicted structures of *Pc*INP. Structures are colored according to the local distance difference test (pLDDT) found in the legend. Cysteines are shown as spheres and colored by element.

**Supplementary Table 1.** Constructs used in this study.

| Plasmid | Name in this study | Description |
| --- | --- | --- |
| pET-21(+) | Empty vector control | *E. coli* expression plasmid |
| pICE_Podila_001 | *PcINP* | *PcINP* |
| pICE_Podila_002 | *PcINP*-tagged | *PcINP*_TEV-HA-6xHis |
| pICE_Podila_006 | *PcINP*-ΔSP | *PcINP*_Δsignal_TEV-HA-6xHis |
| pICE_Podila_007 | SUMO-*PcINP*-ΔSP | 6xHis_SUMO_*PcINP*_Δsignal_TEV-HA |
| pICE_Podila_008 | SUMO-*PcINP*-ΔSPΔN | 6xHis_SUMO_*PcINP*_Δsignal_ΔN-term_TEV-HA |
| pICE_Podila_009 | *PcINP*-ΔSPΔN | *PcINP*_Δsignal_ΔN-term_TEV-HA-6xHis |

**Supplementary Fig 3.**
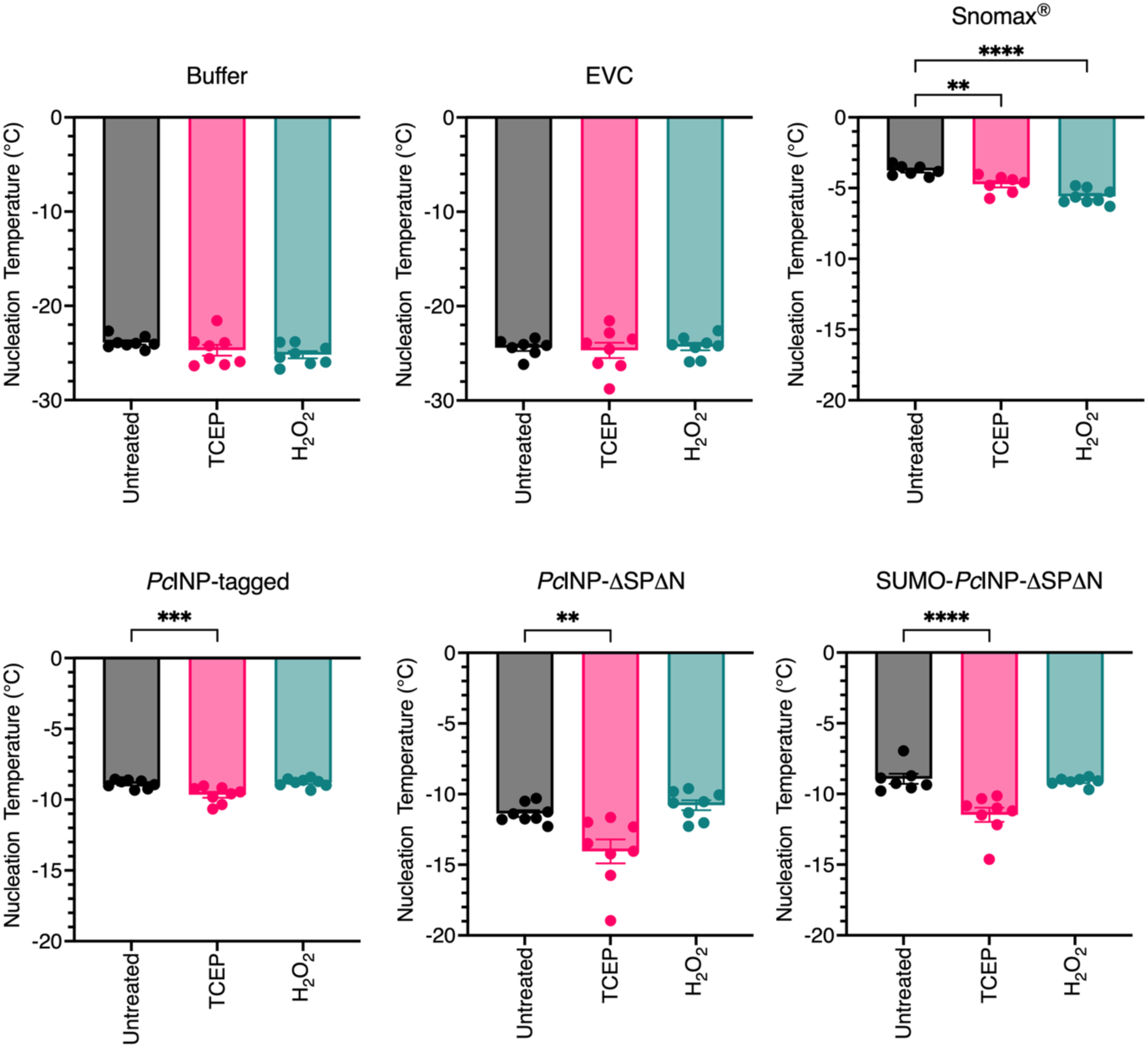
Impact of cysteine redox state of *Pc*INP on ice nucleation activity. Samples were treated with tris(2-carboxyethyl)phosphine (TCEP) or hydrogen peroxide (H_2_O_2_) for 1 hr before ice nucleation activity was measured. Data are presented as mean ± s.e.m. Statistical analysis was performed using a one-way ANOVA.

**Supplementary Fig 4.**
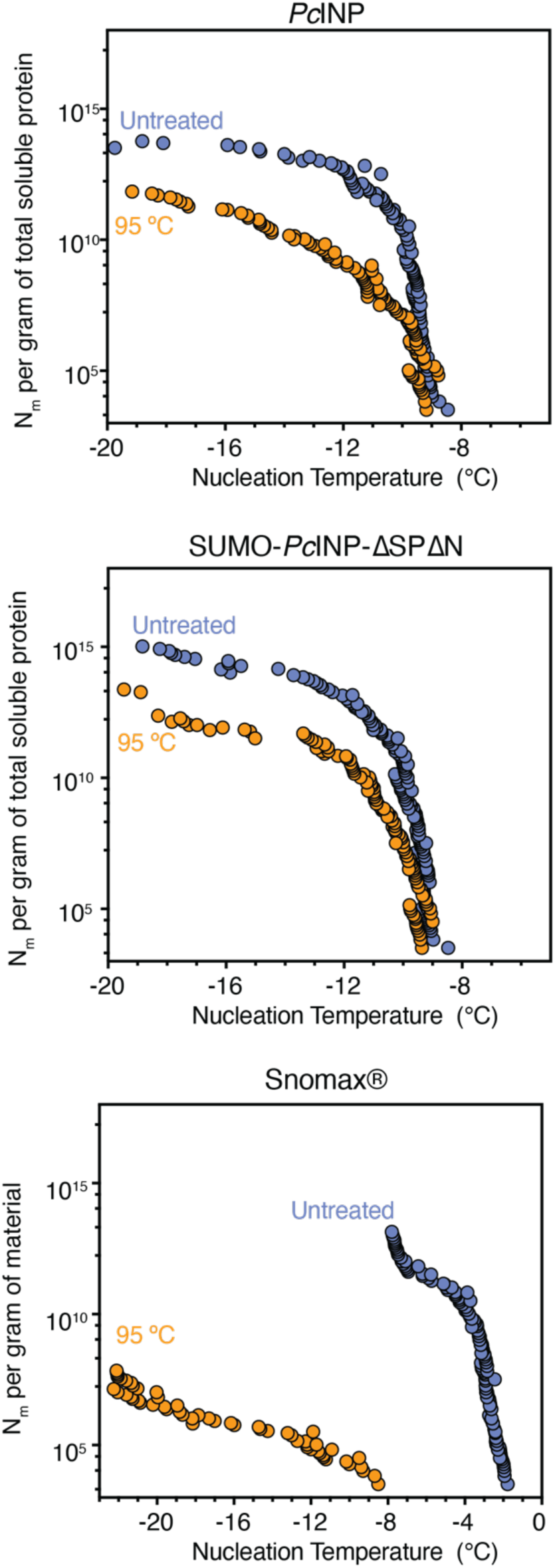
Impact of temperature on ice nucleation activity. Number of ice nuclei (N_m_) per gram of total protein measured for soluble protein lysates of *Pc*INP and SUMO-*Pc*INP-ΔSPΔN, and N_m_ per gram of dry weight of Snomax^®^ with and without treatment at 95 °C before dilution.

**Supplementary Fig 5.**
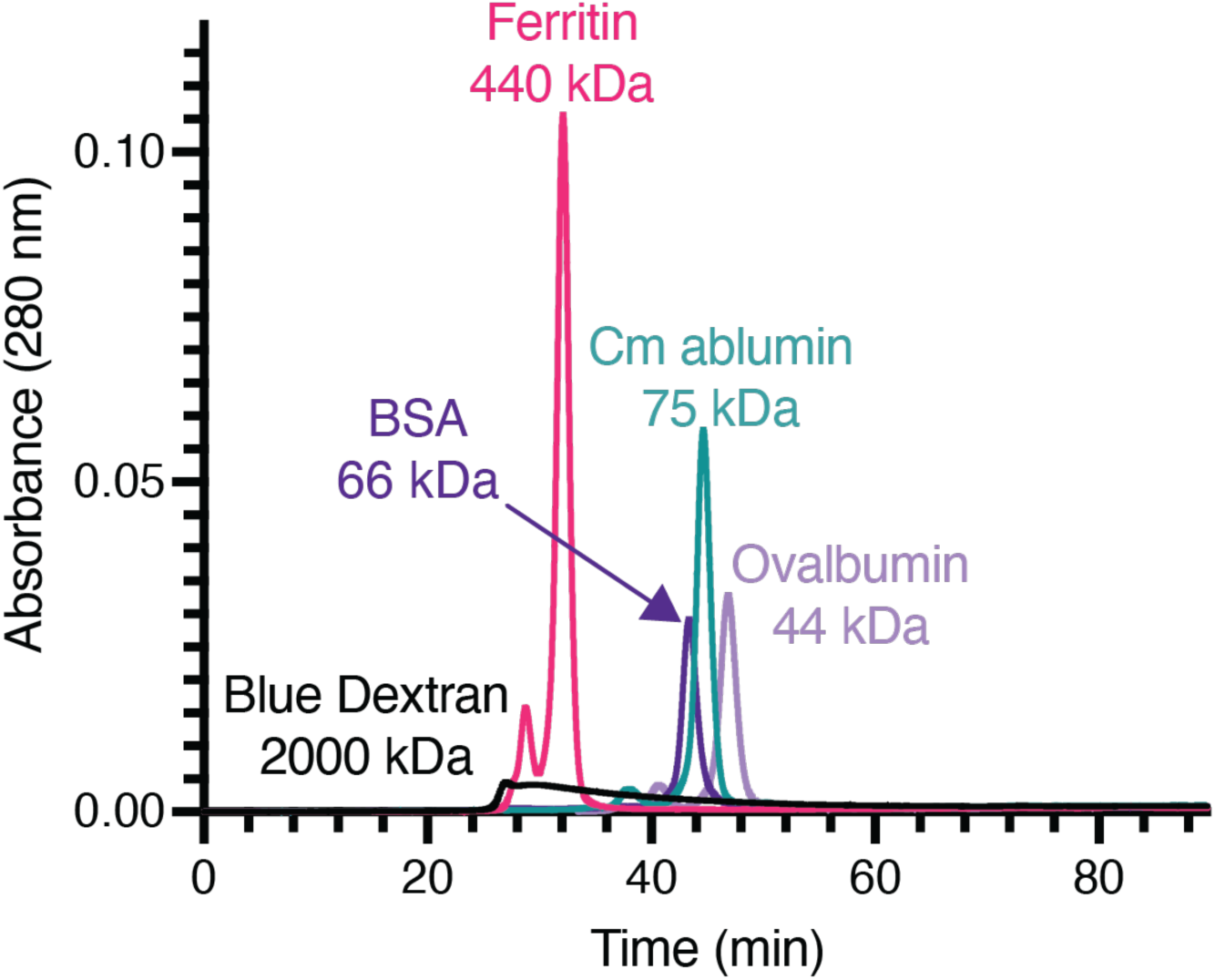
SEC-MALS of protein standards. Protein standards ranging from 44 kDa to 440 kDa were analyzed by SEC-MALS to determine retention time range for monomer and oligomerized proteins. Blue dextran was used to determine the void volume of the column.

